# Psychedelic Hormesis: LSD Activates Adaptive Stress Transcriptional Programs in the Prefrontal Cortex

**DOI:** 10.64898/2026.09.11.751070

**Authors:** Aurora Savino, Carla Liaci, Giorgio Roberto Merlo, Lucia Prandi, Lidia Avalle, Valeria Poli

## Abstract

**Background:** Classic psychedelics are serotonergic agents increasingly recognized for their ability to produce rapid and long-lasting therapeutic effects in several neuropsychiatric conditions, yet the molecular mechanisms that translate acute serotonergic perturbation into long-term brain adaptation remain poorly understood. Psychedelic action is commonly attributed to enhanced neuronal plasticity, but emerging evidence suggests broader engagement of stress-responsive and homeostatic processes across neural and non-neural cell types.

**Methods:** To define the temporal structure of psychedelic-induced transcriptional responses, we profiled gene expression in the prefrontal cortex of mice at early and delayed time points following a single administration of lysergic acid diethylamide (LSD). Differentially expressed genes were then contextualized through pathway-level and cross-dataset comparisons with established models of adaptive and chronic stress.

**Results:** LSD induced a coherent, temporally organized transcriptional program extending beyond immediate neuronal activation. Early responses reflected adaptive metabolic and stress-related signaling, whereas later ones involved circadian and neuroendocrine regulation. A persistent transcriptional core spanning both time points indicated sustained regulation of metabolic, circadian, and stress-responsive pathways. Comparative analyses showed that, although LSD-induced transcriptional profile shares features with general stress responses, its preferentially aligns with adaptive, hormetic stress paradigms rather than maladaptive chronic stress.

**Conclusions:** These findings indicate that in the prefrontal cortex LSD activates adaptive stress transcriptional programs consistent with hormetic adaptation, providing a molecular framework to interpret the enduring effects of psychedelics beyond synaptic plasticity.

## Background

Classic psychedelics such as lysergic acid diethylamide (LSD), psilocybin, and N,N-dimethyltryptamine exert strong serotonergic effects that acutely alter perception, cognition, and affect [1, 2]. After decades of limited clinical investigation, these agents have re-emerged as promising therapeutics for mood, anxiety, trauma-related, and substance use disorders [3–5]. In contrast to conventional serotonergic antidepressants, whose efficacy typically depends on prolonged administration and is often limited in treatment-resistant populations, psychedelics can produce rapid and sustained clinical improvements following one or a few administrations [3, 6–8]. This unusual feature has motivated intense investigation into the molecular mechanisms that translate acute drug exposure into long-lasting behavioral change.

Pharmacologically, classic psychedelics primarily engage the serotonin 5-HT_2A_ receptor, which is highly expressed in cortical association areas and necessary for the molecular, hallucinogenic and behavioral effects of psychedelics [4, 9–18]. Receptor activation elicits transient induction of immediate-early genes (IEGs), increases in cortical excitability, and reconfiguration of brain functional connectivity toward higher entropy [19, 20]. While these acute circuit-level effects are thought to underlie the subjective psychedelic experience, they cannot fully account for the persistence of therapeutic benefits, implicating longer-lasting molecular and structural adaptations. Consistent with this view, extensive preclinical work demonstrates that psychedelics robustly promote structural and functional neuroplasticity [21–25]. LSD and psilocybin increase dendritic spine density, enhance synaptogenesis, and stimulate neurite outgrowth both *in vitro* and *in vivo* [18, 26–29]. These effects depend on coordinated activation of neurotrophic and translational signaling pathways, including BDNF-TrkB, mTOR and ERK, and persist well beyond the period of acute receptor engagement [30–32]. Recent studies further reveal that psychedelic-induced plasticity is cell-type- and circuit-specific, involving discrete populations of cortical pyramidal neurons whose activation is sufficient to recapitulate antidepressant- and anxiolytic-like behaviors [24, 33, 34]. Yet, the transcriptional programs coordinating these durable changes remain incompletely defined. While early transcriptomic studies emphasized acute induction of IEGs such as *Fos*, *Egr1/2*, *Arc*, and *Nr4a* family members shortly after psychedelic exposure, reflecting rapid neuronal activation [11, 34, 35] more recent transcriptomic and epigenomic analyses demonstrate that psychedelics engage broader and more persistent gene-expression programs [36–43]. Notably, these responses also involve non-neuronal cell types such as glial and immune cells, suggesting engagement of multi-cellular adaptive programs [36–43]. Beyond providing mechanistic insight into the molecular effects of psychedelic exposure, transcriptomic profiling offers a quantitative framework for situating these effects within a broader landscape of biological states. By summarizing coordinated gene-expression changes across pathways and cellular programs, transcriptional signatures can be treated as high-dimensional readouts that enable systematic comparison between pharmacological perturbations and other physiological or pathological conditions, thereby facilitating an integrative, cross-condition interpretation of their biological significance [44–49].

Psychedelics’ enduring effects on anxiety- and stress-related behaviors [7, 50–56]. are difficult to reconcile with transient changes in neuronal excitability or synaptic structure alone, suggesting instead a shift in the organism’s ability to respond to stressors, i.e. an increase in stress resilience [57]. Stress resilience is the unifying emerging effect of coordinated molecular programs that enable tissues and neural circuits to absorb, adapt to, and recover from acute perturbations while maintaining functional stability [57–59]. Importantly, such adaptive responses are inherently non-linear. While transient and moderate stressors engage adaptive programs that support synaptic plasticity, metabolic efficiency, and learning, sustained or excessive stress leads to functional breakdown [60, 61]. This principle has been extensively characterized in the context of glucocorticoid signaling, where activation of high-affinity, low-capacity mineralocorticoid receptors promotes neuroplasticity and energy utilization, whereas prolonged engagement of low-affinity, high-capacity glucocorticoid receptors disrupts plasticity and mediates stress-related toxicity [60, 61]. The biological framework of hormesis provides a formalization of this non-linear adaptive logic. Hormesis describes a biphasic dose-response relationship in which transient, moderate perturbations activate adaptive stress-response programs that enhance cellular resilience and plasticity while avoiding the deleterious consequences of chronic stress exposure [62, 63]. A wide range of hormetic stimuli, including physical exercise, intermittent fasting, metabolic challenge, and environmental enrichment, are known to induce overlapping transcriptional programs involving mitochondrial biogenesis, redox regulation, autophagy, metabolic rewiring, and neurotrophic signaling/neuroplasticity [63–65].

Here, we characterize the transcriptional response to LSD in the prefrontal cortex of mice at both early (90 minutes) and delayed (24 hours) timepoints, revealing a coherent and temporally structured transcriptional program that extends beyond immediate neuronal activation and strongly overlaps with established hormetic and adaptive stress-response signatures. Our work provides a mechanistic bridge between acute serotonergic perturbation, conserved adaptive stress responses, and sustained neural plasticity, highlighting transcriptional hormetic programs as a central substrate of psychedelics’ enduring effects.

## Methods

The primary objective of this study was to characterize the molecular and transcriptional changes induced by LSD in the prefrontal cortex over short- and intermediate-term timescales, and to explore the biological mechanisms underlying these responses in order to better understand their potential functional consequences, therapeutic relevance, and associated risks.

We first focused on defining the global structure and temporal dynamics of the LSD-induced transcriptional response by integrating differential expression and pathway-level analyses at early (90 min) and delayed (24 h) time points. This discovery-driven approach was designed to identify coherent biological programs activated by LSD, rather than to test predefined hypotheses about specific pathways or phenotypic outcomes.

During this initial characterization, we observed that several LSD-responsive transcriptional programs were enriched for processes classically associated with cellular and systemic stress responses, including metabolic adaptation, neuroendocrine signaling, and regulatory feedback pathways. This observation motivated a secondary, data-informed analytical step in which LSD-induced signatures were contextualized relative to external transcriptomic datasets representing diverse forms of physiological hormetic stress and maladaptive chronic stress.

This analytical strategy allowed us to (i) describe the core molecular effects of LSD in the prefrontal cortex, (ii) generate mechanistic hypotheses about the nature of these responses, and (iii) evaluate their broader biological context in relation to known stress paradigms.

### Mice housing and treatment

8 weeks old male C57BL/6 mice were raised and maintained in the specific pathogen-free transgenic unit of the Molecular Biotechnology Center (University of Turin) under a 12-h light/dark cycle and provided food and water ad libitum. Procedures were conducted in conformity with national and international laws and policies as approved by the Faculty Ethical Committee and the Italian Ministry of Health.

Mice were injected intraperitoneally with either saline (0.9% NaCl) vehicle or LSD (500 ug/kg) and euthanized after 90 minutes or 24 h through cervical dislocation. The animals were monitored throughout the treatment period, and the treatment was considered effective based on the presence of head-twitch response in the 15 to 30 minutes timeframe after treatment [66].

### RNA-sequencing

Brains were removed and sectioned on a vibratome. The range of sections used for analysis corresponds to coronal sections 25-43 of the reference Allen Brain Atlas (atlas.brain-map.org). The prefrontal cortex areas were isolated and selected following published protocols [67]. Total RNA was extracted using QIAzol reagent and RNeasy Lipid Tissue kit (Qiagen) according to manufacturer instructions. RNA Integrity was quantified with the TapeStation (Agilent) automated electrophoresis system, ensuring a RIN > 8 for libraries preparation. The Zymo-Seq RiboFree Total RNA Library Kit (Zymo Research) was employed for library preparation and a minimum of 30 million Paired-end 150bp reads per sample were sequenced. For data pre-processing, we employed an adapted version of the nf-core RNAseq pipeline v2.1.0 (https://zenodo.org/records/17909656)[68]. Specifically, we checked the quality of the sequencing with FastQC v0.11.9 (https://www.bioinformatics.babraham.ac.uk/projects/fastqc/), aligned the reads to the Grcm39 mouse genome with STAR v2.6.1d [69], and counted the reads aligned to each gene with featureCounts v2.0.1 [70].

### RNA-seq preprocessing and gene annotation

Raw count matrices were processed in R. Ensembl gene identifiers were mapped to official *Mus musculus* gene symbols using biomaRt [71]. When multiple Ensembl IDs mapped to the same gene symbol, only the entry with the highest mean expression across samples was retained, thereby avoiding duplicated rows and favoring the most consistently expressed transcript. Genes without an unambiguous symbol assignment were discarded.

Lowly expressed genes were removed by retaining only those with at least 10 counts in more than two samples. For exploratory analyses, count matrices were normalized to reads per million (RPM) and log-transformed with an offset of 1. All downstream differential expression models were fitted on raw counts. To characterize the global structure of the LSD transcriptomic response, Principal Component Analysis (PCA) was performed on log_2_(RPM + 1) values using the FactoMineR package [72].

### Differential expression analysis

Differential expression (DE) between LSD-treated and saline-treated animals was assessed separately at 90 min and 24 h using DESeq2 [73]. For each contrast, DESeq2 models were fitted on raw count matrices with treatment as the main design factor. Wald tests were used to obtain log2 fold-changes (log_2_FC) and nominal p-values for each gene. Unless otherwise specified, genes with Benjamini-Hochberg False Discovery Rate (FDR)-adjusted p < 0.05 were considered significantly differentially expressed and used to define LSD-specific up- and down-regulated gene sets.

### Over-representation analysis of GO Biological Processes

To link LSD-induced gene expression changes to biological processes, we performed over-representation analysis for significantly upregulated and downregulated genes at each LSD time point. For each gene set (up and down) and time point, enrichGO from clusterProfiler [74] was applied using the org.Mm.eg.db annotation package and the Gene Ontology Biological Process (BP) ontology. The universe was defined as all genes tested by DESeq2 for the corresponding contrast. Multiple testing correction was performed using the Benjamini-Hochberg method, and enriched categories were retained using nominal p < 0.01 and FDR < 0.05. Dot plots were used to visualize the most enriched categories.

### Public dataset retrieval and inclusion criteria

To contextualize LSD-induced transcriptional changes, we re-analyzed a curated set of publicly available rodent whole brain and cerebral cortex transcriptomic datasets spanning fasting and caloric restriction, ischemic preconditioning, exercise, and chronic stress paradigms (Suppl. Table 1). Datasets, derived from both RNA-seq and microarray platforms, were retrieved from the Gene Expression Omnibus [75] (GEO, https://www.ncbi.nlm.nih.gov/geo/) via GEOquery [76]. For each series, we selected brain or cortical samples and contrasts upon a hormetic or chronic stress intervention, defined as follows:

–Dietary hormesis: acute or intermittent fasting, ketogenic or calorie-restricted diet, time-restricted feeding versus ad libitum or control diet, including age- or sex-stratified comparisons where available.

–Ischemic / hypoxic / exercise hormesis: brief ischemia or ischemic preconditioning versus sham, hypoxia versus normoxia, and treadmill exercise combined with an MPTP lesion.

–Chronic stress: chronic restraint or multimodal stress, chronic mild stress, chronic social defeat stress, and other chronic stress paradigms.

Within each series, contrasts were defined to compare the hormetic or stress condition against a matched control within a homogeneous subset of samples (same tissue, genotype, sex and, where possible, age).

### Gene identifier mapping and symbol harmonization

For RNA-seq datasets with Ensembl identifiers, Ensembl IDs were mapped to official gene symbols using biomaRt and the appropriate *Mus musculus* or *Rattus norvegicus* Ensembl mart. For microarray datasets, probe identifiers were mapped to gene symbols using the relevant GEO platform annotation. When multiple IDs mapped to the same gene symbol, only the ID with the highest mean expression across samples was retained. Genes without an unambiguous symbol assignment were removed.

### Preprocessing of RNA-seq datasets

For RNA-seq datasets provided as raw counts, we worked directly on count matrices after gene-ID harmonization. DE analyses were then performed using DESeq2 [73]. For datasets provided as FPKM values, we used limma [77] on log2-transformed values to compute differential expression. In all cases, contrasts were defined to reflect the intervention (hormesis or stress) versus its matched control.

### Preprocessing of microarray datasets

For microarray datasets, expression values were obtained from the GEO series matrix, mapped to gene symbols using the corresponding platform annotation, and transformed as log_2_, with an offset of 1, where needed. Linear models were fitted with limma [77].

### Quantifying overlap between LSD signatures and hormetic or chronic stress datasets

To quantify the similarity between LSD-induced transcriptional signatures and external hormesis or chronic stress signatures, we systematically tested enrichment of LSD DEGs in each external dataset. Within each dataset, DE genes were defined as those with nominal p < 0.05 to increase sensitivity in overlap testing, while FDR thresholding was applied at the level of overlap p-values.

For each block of datasets (hormesis or chronic stress), we iterated over both LSD reference signatures (90 min and 24 h) and all target datasets belonging to that block. For a given LSD reference and target dataset, the background gene universe was defined as the intersection of genes present in both DE tables; comparisons with fewer than 50 shared genes were discarded. We then performed a one-sided Fisher’s exact test to assess whether the overlap between LSD DEGs and target DEGs was enriched beyond chance (odds ratio OR > 1). Within each block, nominal overlap p-values were adjusted using the Benjamini-Hochberg method.

### Directional overlap analyses

The same framework was extended to incorporate directionality of regulation. Within each dataset, we defined up-regulated genes as those with logFC > 0 and nominal p < 0.05, and down-regulated genes as those with logFC < 0 and nominal p < 0.05. For each LSD reference and external dataset, we evaluated four directional overlaps: LSD_UP vs DATASET_UP, LSD_DOWN vs DATASET_ DOWN, LSD_ UP vs DATASET_ DOWN, and LSD_ DOWN vs DATASET_UP. For each combination, a one-sided Fisher’s exact test (OR > 1) was performed using the same background gene universe and FDR correction. This analysis allowed us to distinguish concordant (same-direction) from discordant (opposite-direction) overlaps between LSD and external signatures.

### Meta-analytic summaries of LSD DEG enrichment

To summarize enrichment across multiple datasets within each block, we computed meta-analytic Fisher combined p-values [78] using the metap package (10.32614/CRAN.package.metap). For each block and LSD reference (90 min and 24 h), we combined the set of nominal overlap p-values across all datasets into a single meta-analytic statistic reflecting the overall strength of evidence for overlap between LSD signatures and that category of interventions. In parallel, we computed the average log odds ratio (mean log OR) across datasets as a complementary measure of effect size.

### Direct comparison of hormetic versus chronic stress overlap with LSD signatures

To directly compare how strongly LSD signatures aligned with hormetic versus chronic stress signatures, we employed three alternative strategies:

– We tested whether hormesis datasets tended to show stronger overlap with LSD signatures than chronic stress datasets by performing one-sided Wilcoxon rank-sum tests on log_2_OR values.

– We compared the overall significance of the overlap obtained through Fisher-aggregated p-values. Because the number of hormesis and chronic stress datasets differed, we implemented a bootstrap-based meta-analytic comparison that explicitly accounts for unequal sample sizes. Let n_s be the number of distinct chronic stress datasets and n_h the number of hormesis datasets. We first computed a meta-analytic Fisher combined p-value for all n_s chronic stress overlaps (p_meta,stress). We then performed B = 5,000 bootstrap iterations. At each iteration, we randomly sampled n_s datasets from the hormesis block without replacement, computed a Fisher combined p-value for that subset, and recorded the result. This procedure generated an empirical distribution of hormesis meta-p-values under the constraint of having the same number of datasets as in the chronic stress block.

– To characterize how the fraction of enriched datasets changes as the significance threshold is relaxed, we constructed a running-threshold. For hormesis and chronic stress blocks separately, we evaluated a grid of p-value thresholds between 0 and 0.2 (200 equally spaced points). For each threshold t, we computed the proportion of LSD-dataset overlaps in that block with nominal Fisher p < t.

### Directional Gene Set Enrichment Analysis (GSEA) against LSD signatures

To quantify both the magnitude and direction of transcriptional responses across heterogeneous datasets, we defined a signed statistic for each gene i:

S_i = sign(log_2_FC_i) × [-log_10_(p_i)],

where log_2_FC_i and p_i are the log_2_ fold-change and nominal p-value from the corresponding differential expression analysis. S_i increases for genes with strong evidence of upregulation and decreases for genes with strong evidence of downregulation. Very small p-values were stabilized by adding 1×10^-300^ before taking the logarithm. For each dataset, genes were ranked in decreasing order of S_i, yielding a genome-wide ordered signature suitable for directional GSEA [79]. This framework enables comparisons across datasets measured on different platforms and tissues and captures directional consistency beyond simple DEG overlap. Query signatures for GSEA were defined for each LSD reference time point (90 min and 24 h) as significantly regulated genes (FDR < 0.05). Among these, genes with log_2_FC > 0 were assigned to the LSD_UP set and those with log_2_FC < 0 to the LSD_DOWN set.

For each hormetic dataset, we ran GSEA via the fgsea R package [80] using the signed ranked statistics as input and the LSD_UP and LSD_DOWN sets as pathways. This yielded Normalized Enrichment Scores (NES), p-values and FDR estimates quantifying the extent to which each hormetic dataset recapitulates the up- or down-regulated LSD programmes. Positive NES indicates alignment with the direction of the LSD gene set, whereas negative NES indicates inverse-alignment.

To summarise both directionality and statistical strength in a single metric, we computed for each GSEA test i:

Z_i = sign(NES_i) × [-log_10_(p_i)],

where NES_i and p_i are the normalized enrichment score and nominal p-value. Z_i is positive when datasets show strongly significant enrichment towards the queried LSD signature. Additionally, for each LSD time point, we counted the number of hormetic datasets with NES > 0 (alignment with the corresponding LSD programme) and NES < 0 (opposite alignment), separately for LSD_UP and LSD_DOWN. We statistically compared the proportion of positive NES between LSD_up and LSD_down with a chi-square test for independence. Significant results indicate asymmetric directional enrichment favoring one LSD programme over the other.

### Pathway-level integration across all datasets

To place LSD, hormetic, and chronic stress responses in a common functional space, we performed pathway enrichment analyses on all available DE results using the same signed-statistic framework. Gene sets were obtained from the MSigDB C5 Biological Process collection via msigdbr (*Mus musculus*) (https://igordot.github.io/msigdbr). For each dataset, the S_i statistic defined above was computed and used to generate a ranked gene list. We then ran a GSEA on these ranked lists, obtaining NES values for each pathway-dataset pair.

### Sparse Partial Least Squares Discriminant Analysis to discriminate hormesis from chronic stress at pathway level

To obtain a low-dimensional discriminant space separating hormetic from chronic stress interventions at the pathway level, we applied sparse Partial Least Squares Discriminant Analysis (sPLS-DA, [81]) to the matrix of NES values. We fitted an sPLS-DA model with 2 components, selecting 20 pathways with non-zero loadings on each component. The training matrix was constructed by selecting all datasets labeled as hormesis or chronic: LSD datasets were withheld from training, ensuring a strict separation between model fitting and LSD projection. This configuration yielded a two-dimensional latent space specifically optimized to discriminate hormesis from chronic stress based on a sparse, interpretable subset of pathways. The model outputs component scores for each dataset and loading vectors that quantify the magnitude and sign of each pathway’s contribution.

To interpret LSD transcriptional responses in the context of hormetic and chronic stress interventions, we projected LSD signatures into the previously learned sPLS-DA space.

To formally quantify whether LSD signatures are closer to hormetic or to chronic stress signatures, we computed pairwise Euclidean distances in the sPLS-DA space between each LSD signature and each hormesis or chronic stress dataset. Centroids were defined as the mean component scores of training datasets within each class. We compared the distributions of LSD-hormesis and LSD-stress distances using Wilcoxon rank-sum tests.

### Stability analysis of LSD proximity to hormesis and chronic stress using sPLS-DA

To assess the robustness of LSD proximity estimates and identify influential training datasets, we performed a leave-one-dataset-out (LODO) analysis across all hormesis and chronic stress datasets.

In each iteration, one training dataset was removed, the sPLS-DA model was refit on the remaining datasets, and LSD projections and distances to class centroids were recomputed. This generated a collection of alternative models, each reflecting the absence of a single training dataset.

For each LSD signature, we computed the frequency with which it was closer to hormesis or stress across all models, providing a measure of classification confidence.

## Results

### Transcriptional profiling of prefrontal cortexes from LSD-treated mice

In order to assess short-term transcriptional responses to LSD, we injected C57BL/6 mice and collected their prefrontal cortexes at 90 minutes and 24 hours (**Fig. 1A**), followed by total RNA extraction and transcriptional profiling by NGS sequencing. The twitching effect described as an early behavioral response to classical psychedelics [82] was assessed by filming the mice in the 15-30 minutes post-injection timeframe and the twitches were counted by an experimenter blind to the treatment (**Suppl. Fig. 1A**).

**Figure 1.**
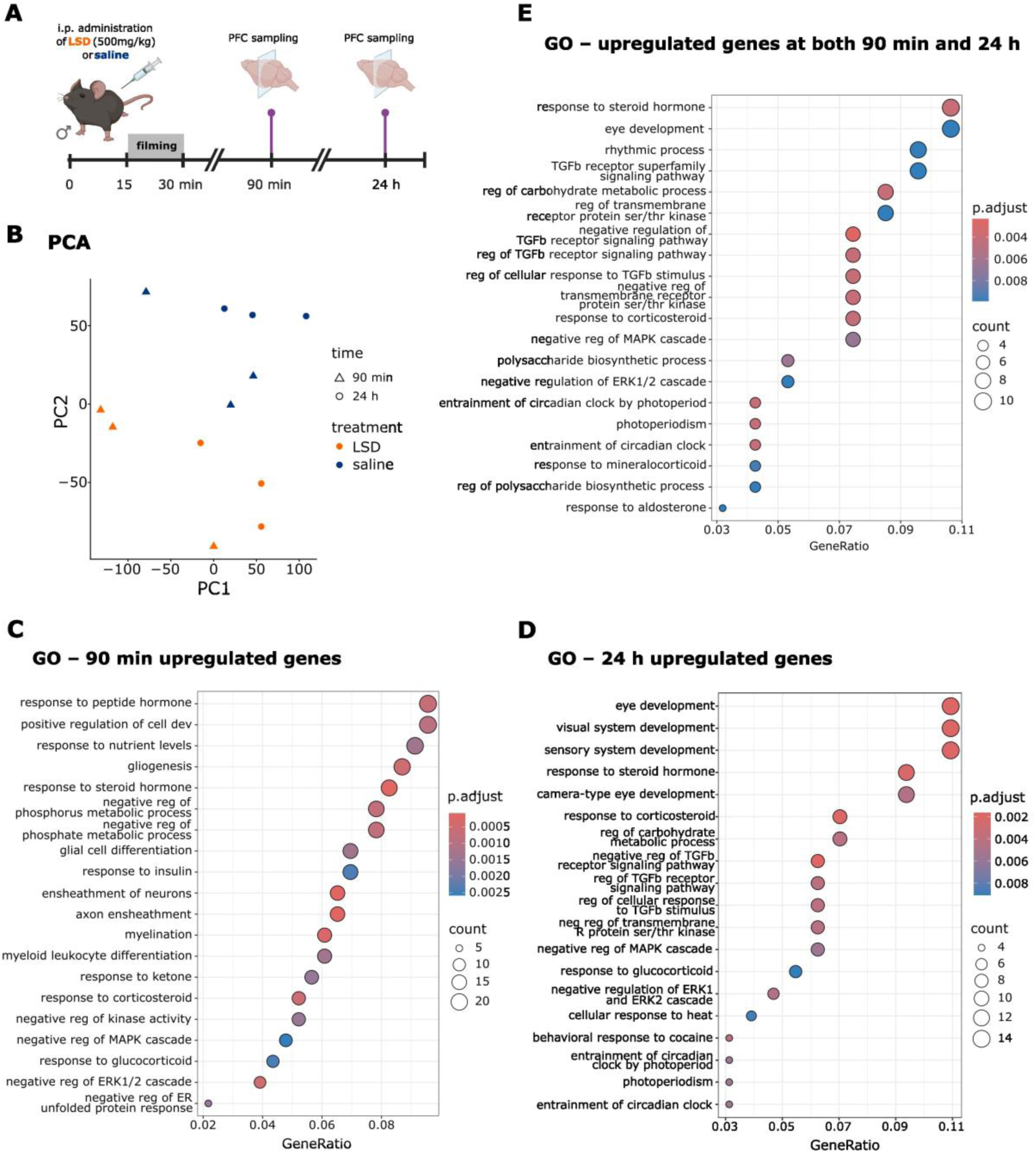
Experimental setup and global transcriptional profiling following acute LSD administration. **A)** Schematic overview of the experimental setup: adult mouse prefrontal cortex samples were collected at 90 min and 24 h after acute intraperitoneal LSD administration. The twitching effect was assessed by filming the mice in the indicated timeframe. **B)** Principal Component Analysis (PCA) of samples labelled by treatment and sampling time. **C–E)** Gene Ontology enrichment analysis of DEGs up-regulated at 90 min (**C**), 24 h (**D**), and at both time points (**E**).

Principal Component Analysis (PCA) of whole-transcriptome profiles revealed a clear separation between LSD-treated and control samples along PC2, indicating that LSD elicits a coherent and large-scale transcriptional shift detectable at both time points (**Fig. 1B**). Differential expression analysis confirmed a robust perturbation, with numerous differentially expressed genes (DEGs): 296 at 90 min and 173 at 24 h with p-adjusted value < 0.05 (**Suppl. Table 2 and Suppl. Fig. 1B**). Interestingly, the differential expression response was biased toward upregulation, with a larger fraction of DEGs showing increased rather than decreased expression at both time points. LSD-responsive genes at both time points include as expected IEGs such as *Fos*, *Egr1/2/3/4*, *Nr4a1/2/3*, *Jun*, *Arc*, *Dusp4/5/6*, *Ier5/5l*, and the neurotrophic factor *Bdnf,* reflecting rapid glutamatergic excitation and activity-dependent transcription (**Suppl. Fig1C**).

The transcriptional program at 90 minutes reveals a rapid, multi-layered response that extends far beyond the canonical neuronal IEG burst classically associated with psychedelics. The dominant features of the 90 min signature arise from coordinated modulation of metabolic, glial, and endothelial pathways (**Suppl. Table 2**). Up-regulated metabolic stress and nutrient-sensing genes (*Txnip*, *Ucp2*, *Pfkfb3*, *Ddit4*, *Sgk1*, *Pdk*, *Gpt2*, *Mat2a*) indicate rapid energy reallocation, redox control, and activation of AMPK/mTOR-linked adaptive programs. Gene Ontology (GO) categories mirror this pattern (**Fig. 1C**): *response to steroid hormone*, *response to nutrient levels*, *response to ketone*, *response to hypoxia*, *generation of precursor metabolites and energy* (**Suppl. Table 3**), consistent with a metabolic stress response. Core circadian clock genes (*Per1*, *Per2*) and clock-linked transcriptional regulators (e.g., *Bhlhe40*, *Klf9*/*Klf10*, *Rasd1*) are up-regulated, consistent with a rapid engagement of circadian transcriptional programs shortly after treatment. In parallel, myelin and OPC-related genes (*Plp1*, *Mbp*, *Mog*, *Cnp*, *Bcas1*, *Ermn*, *Mobp*) indicate over-expression of glial and oligodendrocyte programs, as evidenced by the GO enrichment for *myelination*, *ensheathment of neurons*, *glial cell differentiation*, and *oligodendrocyte differentiation*. These pathways suggest that LSD engages glial structural plasticity, lipid metabolism, and myelin dynamics rather than impacting exclusively neuronal transcription. A third component is the induction of endothelial and vascular homeostasis genes, including *F3 (Tissue Factor)*, *Ccn1*, *Mfsd2a*, *Ddah1*, *Pdgfb*, and *Sdc4*. Finally, the early response includes robust enrichment of negative regulators of the ERK/MAPK cascade (*negative regulation of ERK1/2*, *negative regulation of kinase activity*), consistent with an initial surge in ERK activity.

By 24 hours, the transcriptome no longer shows the early excitatory/metabolic burst observed at 90 minutes. IEGs subside, and the biological program shifts toward circadian, endocrine, and neurovascular remodeling (**Suppl. Table 2)**. A defining feature of the late response is the activation of circadian regulators, including *Per1*, *Per2*, and *Bhlhe40*, accompanied by GO enrichments in *entrainment of circadian clock*, *photoperiodism*, and *rhythmic process* (**Fig. 1D, Suppl. Table 3**). This suggests that a single LSD exposure induces a persistent re-entrainment of the circadian machinery in the prefrontal cortex, consistent with the known serotonergic regulation of CLOCK/PER pathways and documented psychedelic effects on sleep-wake timing [83]. The activation of the endocrine/steroid axis remains a prominent component also at 24 hours. Several early metabolic-hormonal regulators (e.g., *Pfkfb3*, *Ptgs2*, *Sdc4*, *Ak4*, *Sgk1*) remain elevated, and GO categories such as *response to corticosteroid*, *response to glucocorticoid*, and *response to steroid hormone* continue to be enriched, indicating a stable neuroendocrine recalibration. The 24 h profile also incorporates growth and trophic factors, including *Bdnf* and *Inhba*, as well as anti-inflammatory and homeostatic regulators (*Tsc22d3*, *Nfkbia*, *Sgk1*). These are accompanied by enriched GO categories related to the regulation of TGF-β signaling, including *negative regulation of TGFβ receptor signaling* and *regulation of response to TGFβ stimulus*. This sustained TGF-β modulation likely contributes to glial-neuronal homeostasis, synaptic stabilization, and microglial state transitions [84, 85]. Finally, the 24 h signature includes astroglial and neurovascular interface genes (*Mertk*, *Etnppl*, *Gjb6*, *Frmd6*, *Pxdn*), indicating refinement of astrocyte-endothelial-neuronal communication. GO enrichments referring to sensory/visually related developmental processes likely reflect modules downstream of Wnt/TGF-β rather than literal sensory system activation.

A subset of 99 genes is consistently regulated at both 90 min and 24 h (**Suppl. Table 2**), revealing a stable core program that persists beyond the acute transcriptional wave. Enriched categories such as *entrainment of circadian clock*, *photoperiodism*, and *rhythmic process* (**Fig. 1E**) indicate that circadian modulation is not a transient state but is initiated early and maintained across time. In parallel, terms including *negative regulation of TGFβ receptor signaling* and *TGFβ superfamily signaling pathway* show that modulation of TGF-β signaling is also a sustained feature of the response. Together, these categories suggest that LSD leaves a lasting imprint on the circadian-endocrine interface, a known point of convergence between serotonin signaling, stress hormone dynamics, and glial homeostasis [83, 86–88]. The set includes coherent regulation of pathways such as *regulation of carbohydrate metabolic process*, *glycogen biosynthetic process*, *polysaccharide biosynthetic process*, and *glucan biosynthetic process,* indicating that the strong nutrient-sensing and metabolic-rewiring pattern observed at 90 min does not fully dissipate. Similarly, several endocrine-related categories (*response to steroid hormone*, *response to corticosteroid*, *response to mineralocorticoid*, *response to aldosterone)* remain enriched in the shared signature. Taken together, these categories delineate the persistent core of the LSD response in the prefrontal cortex. This core is defined by sustained circadian regulation, endocrine and glucocorticoid sensitivity, carbohydrate and glycogen metabolic adjustment and modulation of TGF-β signaling.

### Cross-Dataset Comparisons Indicate Broad Overlap with Stress Transcriptomes

The transcriptional programs induced by LSD in the prefrontal cortex described above clearly extend beyond canonical neuronal plasticity, engaging a broad metabolic-endocrine-circadian stress-response axis. This combination strongly resembles the transcriptional architecture triggered by adaptive hormetic stimuli such as fasting, exercise, caloric restriction and ischemic preconditioning - short, controlled physiological challenges that induce metabolic reconfiguration, neurovascular adaptation, circadian entrainment, and resilience-enhancing plasticity [64]. This observation raises the question of whether the LSD signature reflects a beneficial hormetic-like program or else it equally overlaps with the transcriptional consequences of chronic or maladaptive stress, which are characterized by inflammatory signaling, loss of plasticity, dysregulated glucocorticoid responses, and dysregulated circadian rhythms [89–91].

To contextualize LSD-evoked transcriptional states, we assembled a compendium of external datasets spanning hormetic stressors (fasting, caloric restriction, exercise, ischemic preconditioning) and chronic stress paradigms (social defeat, chronic restraint, chronic variable stress), covering 19 datasets across different brain regions and mouse strains (**Suppl. Table 1**). This framework enabled us to ask whether LSD resembles adaptive, transient metabolic stressors or detrimental, prolonged stress exposure. Fisher’s exact tests revealed significant overlap between LSD-regulated genes and DEGs from both hormetic and chronic stress datasets (**Fig. 2A,B**, aggregated p-values < 10^-63^ for both comparisons), suggesting that LSD engages elements of a generic acute stress-responsive transcriptional module. While overlap was detectable for both classes, hormetic paradigms displayed slightly higher median odd ratios (ORs) than chronic stress (**Suppl. Fig. 2A**). However, the overall modest p-values and median differences prompted us to perform more formal and direction-resolved comparisons. To systematically assess alignment strength, we compared the overall significance of the overlap obtained through Fisher-aggregated p-values. Because hormesis and chronic stress blocks included different numbers of datasets, we implemented a bootstrap meta-analytic test to evaluate their relative similarity under equal sample size. (**Suppl. Fig. 2B**). Alternatively, to examine overlap patterns across a continuum of significance cutoffs, we constructed running-threshold curves for the LSD signatures. For hormetic and chronic stress datasets separately, at each p-value threshold, we computed the fraction of datasets exhibiting nominal significance (**Suppl. Fig. 2C**). Comparing these curves provides a threshold-free assessment of whether hormetic interventions systematically produce more frequent or stronger overlaps with LSD than chronic stress paradigms, independent of arbitrary significance cutoffs. None of these analyses revealed a significant difference in the likelihood of LSD-signatures to overlap with hormetic or chronic stress signatures. Notably, Fisher’s tests between hormetic and chronic stress datasets also revealed substantial DEGs’ overlap, suggesting the existence of a shared “generic stress” transcriptional core, making it difficult to distinguish between the two conditions (**Suppl. Fig. 3**).

**Figure 2.**
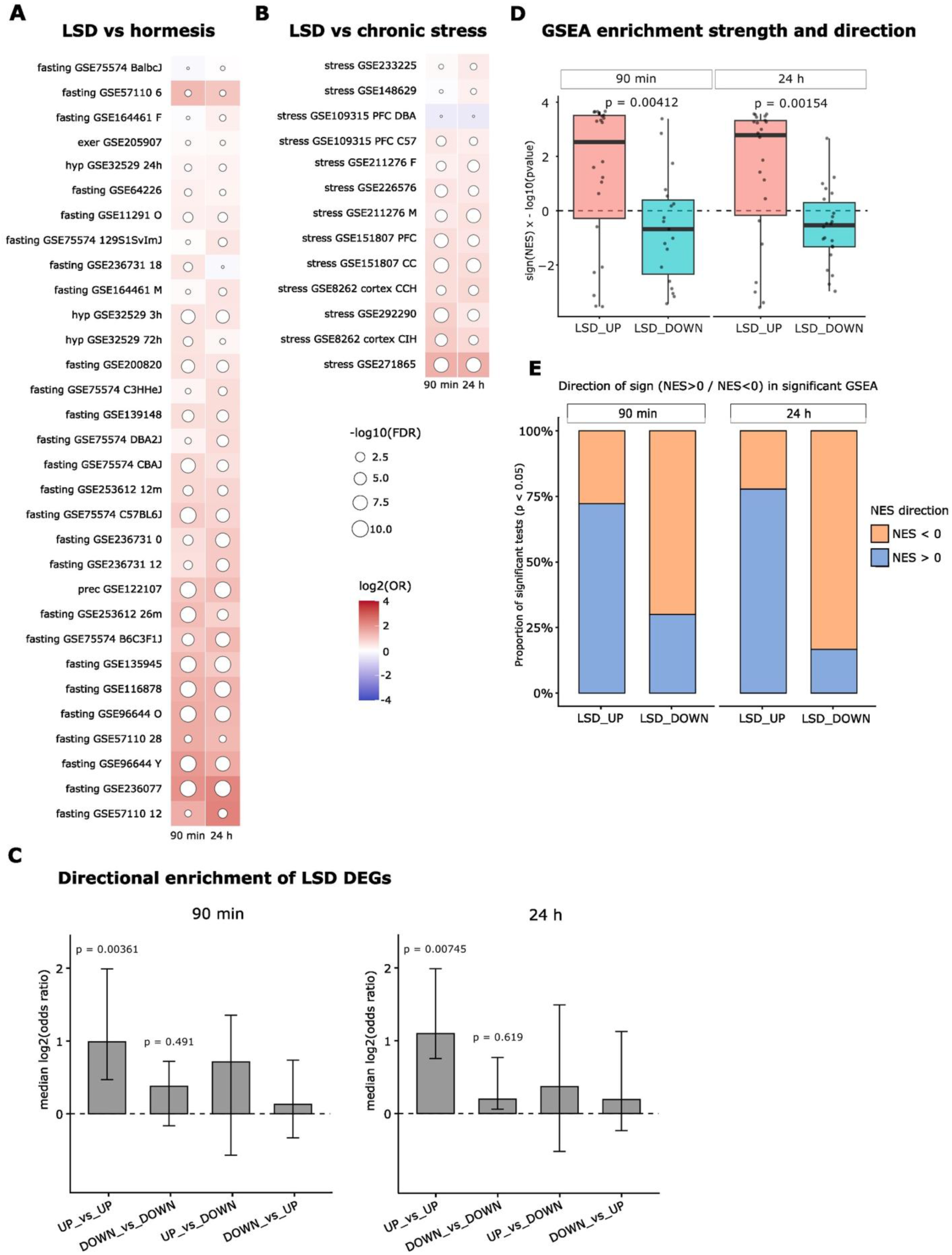
Comparative analysis of LSD transcriptional signatures across hormetic and chronic stress datasets. **A-B)** Overlap between LSD DEGs and external hormesis (**A**) or chronic stress (**B**) datasets, quantified by Fisher’s exact test. Circle size indicates false discovery rate (-log_10_(FDR)), while tile colour indicates the odd ratio (OR). For visualization, -log_10_(FDR) values were capped at 10. **C)** Directional overlap between LSD and hormetic transcriptional signatures. Odds ratios from direction-resolved Fisher’s exact tests comparing LSD-induced DEGs with hormetic stress signatures at 90 min and 24 h post-LSD (Fisher tests across LSD_UP vs DATASET_UP, LSD_DOWN vs DATASET_ DOWN, LSD_ UP vs DATASET_ DOWN, and LSD_ DOWN vs DATASET_UP). We tested whether ORs for UP_vs_UP or DOWN_vs_DOWN Fisher tests were higher than tests on discordant signs (UP_vs_DOWN and DOWN_vs_UP) via a one-sided Wilcoxon test. **D)** Gene set enrichment analysis (GSEA) of LSD_UP and LSD_DOWN signatures ranking genes by differential expression upon hormetic stressors across all hormesis datasets. The boxplots show a signed statistics (-log_10_(p) * sign(NES)) for the GSEA of LSD_UP and _DOWN signatures at 90 min and 24 hours, separately. NES = Normalized Enrichment Score. Significance was assessed via a one-sided Wilcoxon rank-sum test. **E)** Proportion of external datasets resulting in significant positive or negative NES values for LSD_UP and LSD_DOWN gene sets.

### Adaptive Stressors Reproduce the Upregulated Component of the LSD Response

To evaluate not only the presence but also the direction of transcriptional similarity between LSD and hormetic stressors, we first evaluated the directional structure of DEGs overlap between LSD and hormetic stressors using four stratified Fisher tests: LSD_UP vs DATASET_UP, LSD_DOWN vs DATASET_DOWN, LSD_UP vs DATASET_DOWN, and LSD_DOWN vs DATASET_UP (**Suppl. Fig. 4**). Across hormetic datasets, UP_vs_UP produced the highest odds ratios (**Fig. 2C**), indicating a predominant concordance between genes upregulated by LSD and those upregulated by adaptive stressors. In a similar direction-aware comparison between LSD and chronic stress signatures, a coherent sign of differential expression achieved higher odd ratios than opposite signs only for the 24h-post-LSD signature (**Suppl. Fig. 5A**). Overall, LSD_UP vs hormesis_UP odd ratios were higher than LSD_UP vs chronic stress_UP odd ratios at both 90 min and 24 h, while the same did not hold true for different combinations of directions (**Suppl. Fig. 5B**). To capture directionality at the whole-transcriptome level, we ranked genes by a signed statistics increasing for confidently upregulated genes and decreasing for confidently downregulated ones (**see Methods**), producing genome-wide ordered signatures for each external dataset, and enabling cross-platform comparison of directionally consistent patterns. For each LSD reference (90 min, 24 h), the LSD_UP and LSD_DOWN DEG sets were used as queries in a gene set enrichment analysis (GSEA) across all hormesis datasets. To integrate direction and statistical strength, we computed a metric combining the GSEA p-value and the Normalized Enrichment Score (NES) sign. This metric showed a significant difference between LSD_UP and LSD_DOWN gene sets, with higher (on average positive) values for LSD_UP gene sets, indicating a tendency toward a coherent change in gene expression upon LSD and hormetic stress (**Fig. 2D**). Directionality was additionally quantified by counting the number of hormesis datasets with Normalized Enrichment Score (NES) > 0 versus NES < 0 for LSD_UP and LSD_DOWN separately. For both time points, a larger fraction of hormetic datasets showed more NES in the expected direction (NES > 0 for LSD_UP and NES < 0 for LSD_DOWN) than by chance (**Fig. 2E**, chi-square p = 0.078 at 90 min and p = 0.029 at 24 h). Together, these results demonstrate that gene expression changes induced by LSD and hormetic stressors tend to be aligned in direction, expecially considering up-regulated genes.

### LSD Transcriptional States Resemble Hormetic Rather Than Chronic Stress When the Shared Core Is Removed

Because the external datasets spanned different platforms, the number of genes consistently detected across all studies was very small (133 genes). This precluded the integration of the datasets directly on gene-level expression changes or log₂ fold-changes for exploratory analyses (e.g. Principal Component Analysis), as such a restricted feature set would provide a biologically unrepresentative projection of global transcriptional structure. To enable a comprehensive, cross-dataset comparison, we therefore performed PCA on pathway-level NES matrices, which summarize coordinated transcriptional activity across thousands of genes and are robust to differences in gene coverage, dynamic range, and platform-specific effects. This pathway-based PCA provides a unified representation of the major variance components across datasets while avoiding the limitations imposed by the sparse gene intersection. Nevertheless, the PCA did not show clustering of datasets based on the treatment type (**Suppl. Fig. 6**), indicating that unsupervised approaches predominantly capture the common “stress core” shared across conditions and are therefore insufficient to resolve the subtler, class-specific transcriptional differences that distinguish adaptive (hormetic) from detrimental (chronic) stress. This motivated the application of supervised methods capable of extracting the components that best separate the two stress classes.

We therefore performed a Sparse Partial Least Squares Discriminant Analysis (sPLS-DA [81]), a supervised multivariate method that jointly performs dimensionality reduction and classification, learning latent components that maximize the covariance between pathway-level features and predefined class labels. We applied it to pathway NES values, training the model to explicitly discriminate hormetic paradigms from chronic stress models. Projection of LSD samples into this discriminant space revealed a consistent pattern: LSD points localized closer to the hormesis centroid than to the chronic stress centroid (**Fig. 3A**). Quantitatively, Euclidean distances to the hormesis centroid were smaller across the first two components (**Fig. 3B**), indicating that, once the shared stress core is accounted for, LSD aligns more strongly with adaptive hormetic programs than with transcriptional signatures of chronic, detrimental stress. This pattern was mainly driven by distances along component 1, as the distances between LSD signatures (90 min or 24 h post-treatment) and chronic stress signatures were, respectively, 6.4 and 5.4 along component 1, and 1.6 and 1.1 along component 2.

**Figure 3.**
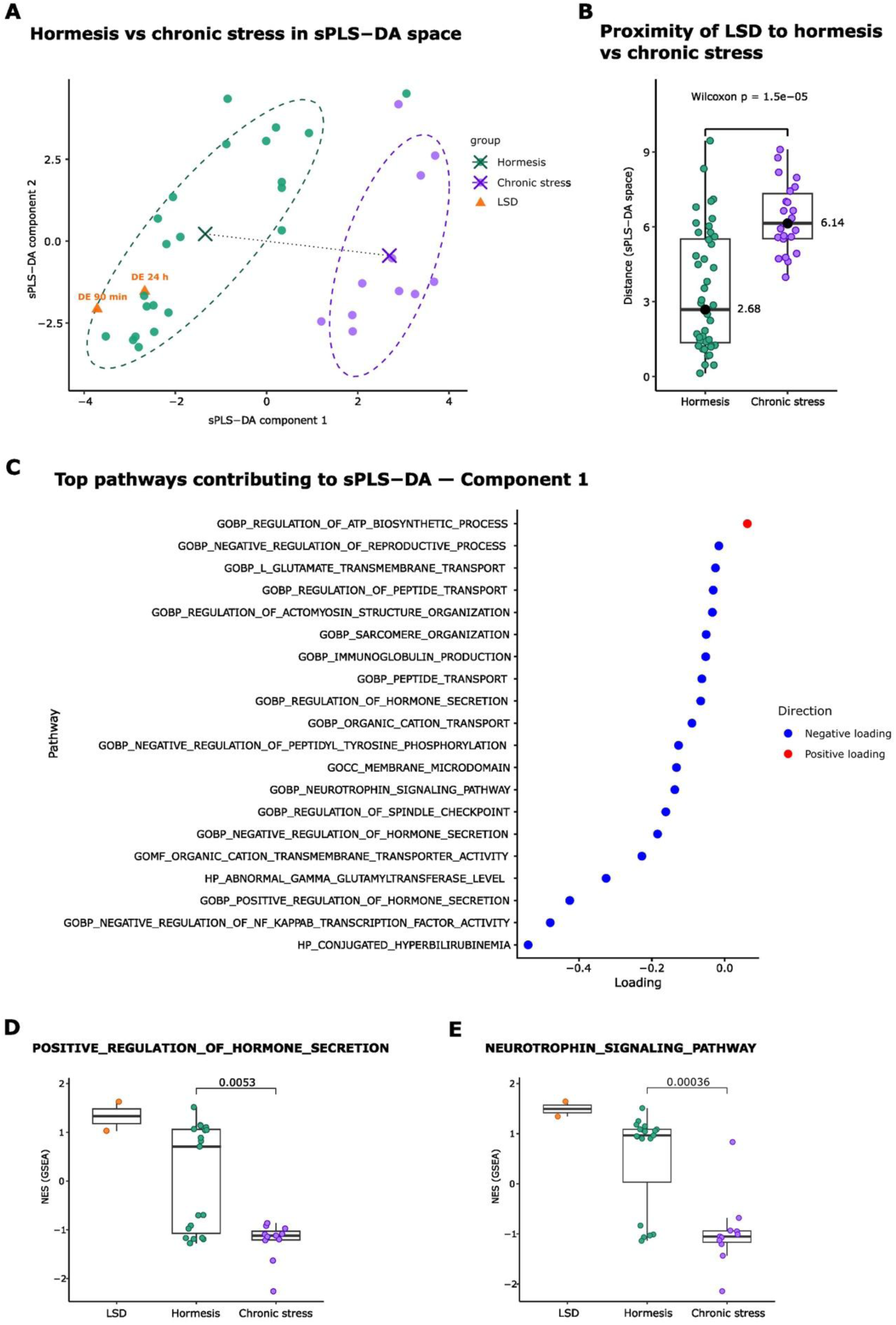
sPLS-DA-based positioning of LSD transcriptional signatures relative to hormetic and chronic stress responses. **A)** sPLS-DA trained on external hormesis and chronic stress datasets, with LSD 90 min and 24 h projected in the latent space. Each point represents a dataset, positioned according to its scores on component 1 and component 2. Ellipses indicate the 68% dispersion contours of the group scores in the sPLS-DA space. **B)** Boxplots of Euclidean distances between LSD samples and the centroids of hormesis or chronic-stress groups. Statistical significance assessed via a two-sided Wilcoxon rank-sum test. **C)** Top contributing pathways and functional categories along sPLS-DA component 1. **D, E)** GSEA Normalized Enrichment Scores of genes ranked by differential expression across treatments – LSD, hormesis and crhonic stress - for the “Positive regulation of hormone secretion” (**D**) and “Neurotrophin signaling pathway” (**E**) categories.

Analysis of pathway loadings (weights assigned to each pathway) revealed that component 1 is dominated by a coherent set of pathways with negative loadings, thus contributing to define the hormetic state, while only a single pathway (*regulation of ATP biosynthetic process*) contributes positively (**Fig. 3C, Suppl. Table 4**). This strong asymmetry indicates that the primary structure captured by component 1 is driven by coordinated variation in pathways activated in the hormetic state. The pathways with negative loadings on component 1 cluster into several biologically related functional themes. A prominent group involves neuroendocrine and hormonal regulation, with pathways such as *positive/negative regulation of hormone secretion* (**Fig. 3D**) and *negative regulation of reproductive process*. A second major theme is signal attenuation and feedback control, exemplified by *negative regulation of NF-κB transcription factor activity* and *negative regulation of peptidyl-tyrosine phosphorylation*. These pathways indicate coordinated dampening of inflammatory and kinase-driven signaling cascades. Several negatively loaded pathways are directly related to neuronal communication and plasticity, including *neurotrophin signaling pathway* (**Fig. 3E**), *membrane microdomain*, *peptide transport*, *regulation of peptide transport*, and *L-glutamate transmembrane transport*. These processes are central to synaptic organization, neurotransmitter handling, and receptor-associated signaling. Other pathways are associated with ionic and molecular transport (*organic cation transport* and *organic cation transmembrane transporter activity*) and structural organization (*sarcomere organization* and *regulation of actomyosin structure organization*), reflecting shared gene modules involved in cytoskeletal organization, membrane dynamics, and intracellular transport, which are also relevant for neuronal structure and signaling. Finally, the presence of Human Phenotype Ontology terms such as *conjugated hyperbilirubinemia* and *abnormal gamma-glutamyltransferase level* most likely reflect metabolic gene modules, rather than tissue-specific pathology. Taken together, these results indicate that component 1 represents a continuum between a state characterized by coordinated regulation of neuroendocrine signaling, synaptic and transport processes, and intracellular signaling restraint (negative direction), and a more limited transcriptional emphasis on ATP biosynthesis and energy regulation (positive direction).

To evaluate the robustness of the sPLS-DA-based positioning of LSD transcriptional signatures relative to hormetic and chronic stress responses, we performed a leave-one-dataset-out (LODO) stability analysis across the training set comprising both hormesis and chronic stress data. In each LODO iteration, a single training dataset was excluded, the sPLS-DA model was refit, and LSD signatures were re-projected into the resulting discriminant space. Distances between LSD signatures and the hormesis and chronic stress centroids were recomputed for each model. Across LODO models, LSD signatures consistently remained closer to the hormesis centroid than to the chronic stress centroid in all iterations (**Suppl. Fig. 7**), indicating that the observed proximity of LSD responses to hormetic programs is not driven by any single external dataset.

## Discussion

Our results demonstrate that LSD elicits a structured and temporally ordered transcriptional response in the prefrontal cortex that extends beyond the canonical IEG burst traditionally associated with psychedelic-induced neuronal activation and plasticity, encompassing metabolic regulation, neuroendocrine signaling, circadian processes, and glial-vascular homeostasis.

Notably, the early 90-minute post-LSD response combines strong IEG induction with coordinated activation of metabolic stress-responsive pathways, glial and oligodendrocyte remodeling programs, and endothelial homeostasis signatures, consistent with an acute adaptive response to energetic or physiological challenge. The convergence of metabolic, glial, and vascular gene programs suggests that LSD transiently shifts the prefrontal cortex into a systemic high-energy, plasticity-permissive state involving multiple non-neuronal compartments. Later, by 24 hours, the transcriptional landscape transitions toward endocrine, circadian, and neurotrophic remodeling. The induction of Per1/Per2 and clock-related processes indicates that serotonergic perturbation produces a reconfiguration of the circadian machinery, consistent with reports of altered sleep-wake timing following psychedelics and with the known serotonergic modulation of circadian oscillators [83]. The persistent enrichment of glucocorticoid- and corticosteroid-responsive pathways points to a neuroendocrine recalibration rather than a transient stress pulse. In parallel, TGF-β- and NF-kB-related regulation and astrocytic structural programsemerge, pointing to slower-acting processes involved in synaptic stabilization, microglial state control, and neurovascular communication.

The gene set coherently modulated at both 90 minutes and 24 hours identifies a lasting “core LSD program”, dominated by circadian entrainment, endocrine sensitivity, modulation of TGF-β signalling, and sustained carbohydrate and glycogen metabolic restructuring. Together, these findings suggest that LSD imparts a coordinated imprint on the circadian-endocrine-metabolic interface of the prefrontal cortex, a functional nexus linked to stress regulation, energy homeostasis, and affective state.

Our cross-dataset comparison framework provides several convergent answers to the key question of whether this core LSD program represents a beneficial, resilience-like adaptation or instead resembles maladaptive chronic stress. When compared to a diverse compendium of hormetic and chronic stress transcriptional signatures, LSD shares a substantial fraction of regulated pathways with both categories. This overlap underscores a key limitation of gene-set intersection-based approaches: shared activation of stress-responsive pathways alone is insufficient to distinguish adaptive from maladaptive biological states. Indeed, substantial overlap was also observed directly between hormetic and chronic stress datasets themselves, indicating the presence of a shared “generic stress” transcriptional core.

To establish whether such similarity reflects concordant or opposing changes, we next examined the directionality of gene regulation. Direction-resolved analyses revealed that the similarity between LSD and hormetic stressors is concordant in sign for the upregulated component of the transcriptional response, while directional concordance was less robust for chronic stress datasets. The considerable overlap of signatures between hormetic and chronic stress datasets, however, complicates interpretation and explains the inability of unsupervised analyses to separate these two stress states. Our supervised pathway-level framework trained to discriminate hormetic from chronic stress paradigms allowed us to overcome this limitation, since LSD signatures consistently localized closer to hormetic than to chronic stress centroids when projected in this discriminant space, indicating that LSD aligns more strongly with adaptive stress responses once the shared stress core is accounted for. This alignment was captured primarily by the first component of the model, dominated by a coherent set of pathways that collectively define the hormetic side of the discriminant axis and converge on several interrelated functional themes. These include regulation of neuroendocrine and hormonal signaling, attenuation of inflammatory and kinase-driven cascades, and processes supporting synaptic function, neurotransmitter handling, and intracellular transport. The coordinated regulation of these pathways points to an integrated transcriptional state characterized by signal modulation, feedback control, and maintenance of neuronal and cellular homeostasis. Moreover, the leave-one-dataset-out analyses demonstrate that the relative proximity of LSD signatures to hormetic datasets is stable, ruling out the possibility that the observed alignment is driven by any single external dataset. These results highlight the importance of supervised, pathway-level analyses for resolving subtle but biologically meaningful differences among correlated transcriptional states.

### Conclusions

Overall, our work reveals that in the prefrontal cortex LSD induces a biphasic, multi-compartment transcriptional program that preferentially aligns with adaptive hormetic stress responses. These findings redefine the molecular architecture of psychedelic action by providing a mechanistic context for how LSD may elicit sustained functional effects, rather than acting solely as a synaptic potentiator. Specifically, LSD appears to reshape brain state by orchestrating an integrated metabolic, endocrine, and glial remodeling program reminiscent of resilience-promoting physiological challenges. This perspective offers a new lens through which to interpret both the therapeutic effects of psychedelics and the potential for synergistic interventions involving stress, metabolism, and circadian biology.

## List of abbreviation

lysergic acid diethylamide (LSD), immediate-early genes (IEGs), differentially expressed genes (DEGs), Differential expression (DE), Principal Component Analysis (PCA), Gene Set Enrichment Analysis (GSEA), Gene Expression Omnibus (GEO), sparse Partial Least Squares Discriminant Analysis (sPLS-DA), False Discovery Rate (FDR), Normalized Enrichment Scores (NES), leave-one-dataset-out (LODO).

## Declarations

– **Ethics approval and consent to participate:** Experimental procedures involving animals were conducted in conformity with national and international laws and policies as approved by the Faculty Ethical Committee and the Italian Ministry of Health.
– **Consent for publication:** Not applicable.
– **Availability of data and materials:** The datasets generated during the current study will be available in the Gene Expression Omnibus repository upon publication. The code to reproduce the analyses is deposited on Github https://github.com/AuroraSavino90/LSD_hormesis
– **Competing interests:** The authors declare that they have no competing interests
– **Funding:** Research reported in this publication was supported by an Impact Initiative Grant of Zymo Research Corporation, by the Italian Ministry of University and Research (MUR PRIN to V.P.), the National Center for Gene Therapy and Drugs based on RNA Technology, Spoke2, PNRR M4C2-Investimento 1.4-CN00000041-VP, and the Truus and Gerrit van Riemsdijk Foundation, Liechtenstein. The content is solely the responsibility of the authors.
– **Authors’ contributions:** A.S., L.A. and V.P. conceived the project, interpreted the data and wrote the manuscript, V.P. coordinated and supervised the whole project, A.S. performed all computational work, G.R.M. supervised the experimental work and performed *in vivo* treatments, C.L and L.P. collected samples and extracted RNA, C.L. and L.A. performed library preparation.

## Acknowledgements

We thank Maddalena Arigoni for her support in libraries preparation and quality checks.

