## Supplementary material for "Psychedelic Hormesis: LSD Activates Adaptive Stress Transcriptional Programs in the Prefrontal Cortex": Suppl info

### SUPPLEMENTARY INFORMATIONS

Supplementary Figure 1……………………………………………………………………………..p.2

Supplementary Figure 2……………………………………………………………………………..p.3

Supplementary Figure 3……………………………………………………………………………..p.4

Supplementary Figure 4……………………………………………………………………………..p.5

Supplementary Figure 5……………………………………………………………………………..p.6

Supplementary Figure 6……………………………………………………………………………..p.7

Supplementary Figure 7……………………………………………………………………………..p.7

Supplementary Tables………………………………………………………………………………p.8

### Supplementary figures

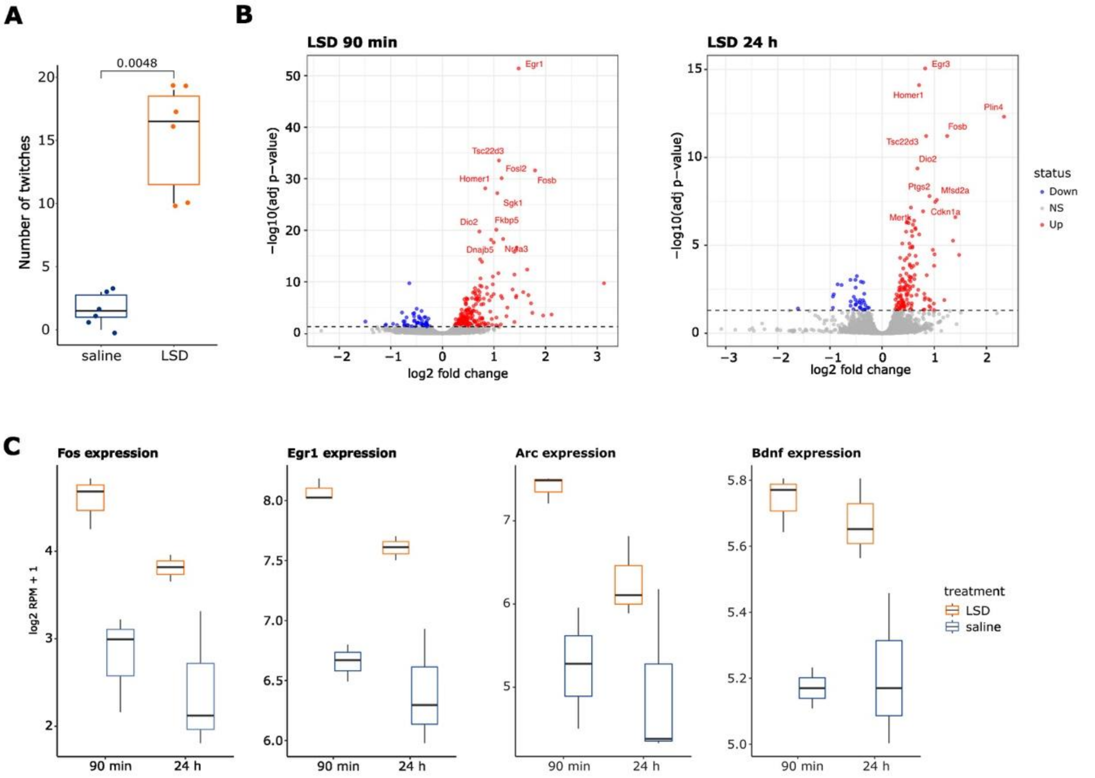


**Supplementary Figure 1. LSD elicits rapid behavioral activation and a robust, time-resolved transcriptional response in the prefrontal cortex. (A)** Boxplot showing the number of head-twitch responses recorded in the 15-30 minutes time frame following acute LSD administration, compared to vehicle controls. Statistical significance was assessed using a two-sided Wilcoxon rank-sum test. **(B)** Volcano plots showing differential gene expression in the prefrontal cortex at 90 min (left) and 24 h (right) after LSD administration versus controls. The x-axis indicates $\log_2$ fold-change, and the y-axis the -$\log_{10}$ adjusted p-value. The top 10 significantly regulated genes are highlighted. **(C)** Boxplots showing normalized expression levels of canonical immediate-early genes (*Fos*, *Egr1*, *Arc*) and the neuroplasticty gene *Bdnf* across experimental conditions and time points.

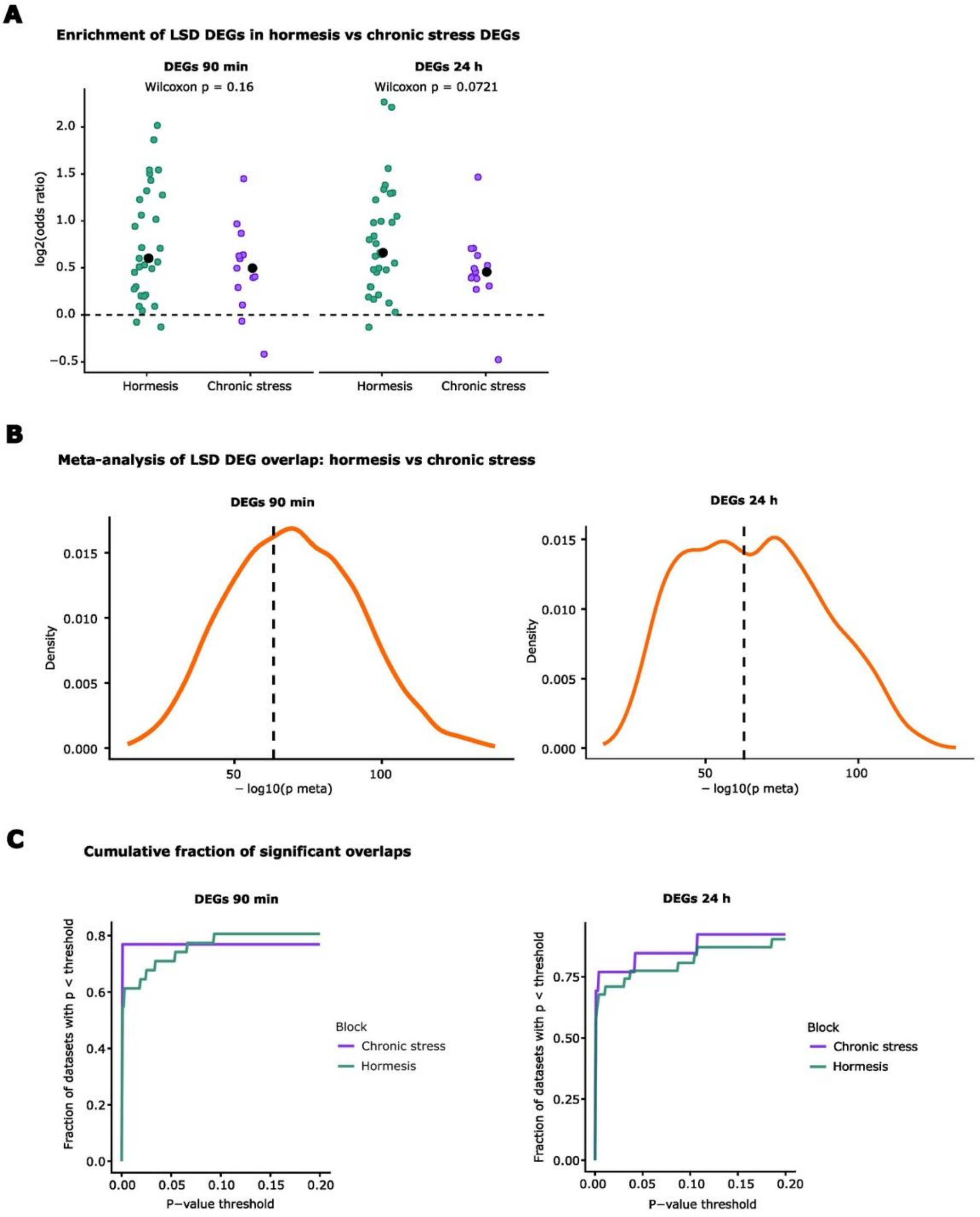


**Supplementary Figure 2. Overlap-based similarity between LSD and stress paradigms. A)** Boxplots showing odds ratios (ORs) from Fisher's exact tests quantifying the overlap between LSD-regulated genes and genes differentially expressed in hormetic or chronic stress datasets. Each point represents a single external dataset. Significance was tested via a one-sided Wilcoxon rank-sum test. **B)** Results of a bootstrap meta-analytic test comparing the overall significance of LSD overlap with hormetic versus chronic stress datasets. At each iteration, equal-sized subsets of datasets were sampled from each class, Fisher-aggregated p-values were computed, and their relative significance was compared. **C)** Running-threshold curves showing, for increasing p-value cutoffs, the fraction of hormetic or chronic stress datasets exhibiting nominally significant overlap with LSD transcriptional signatures.

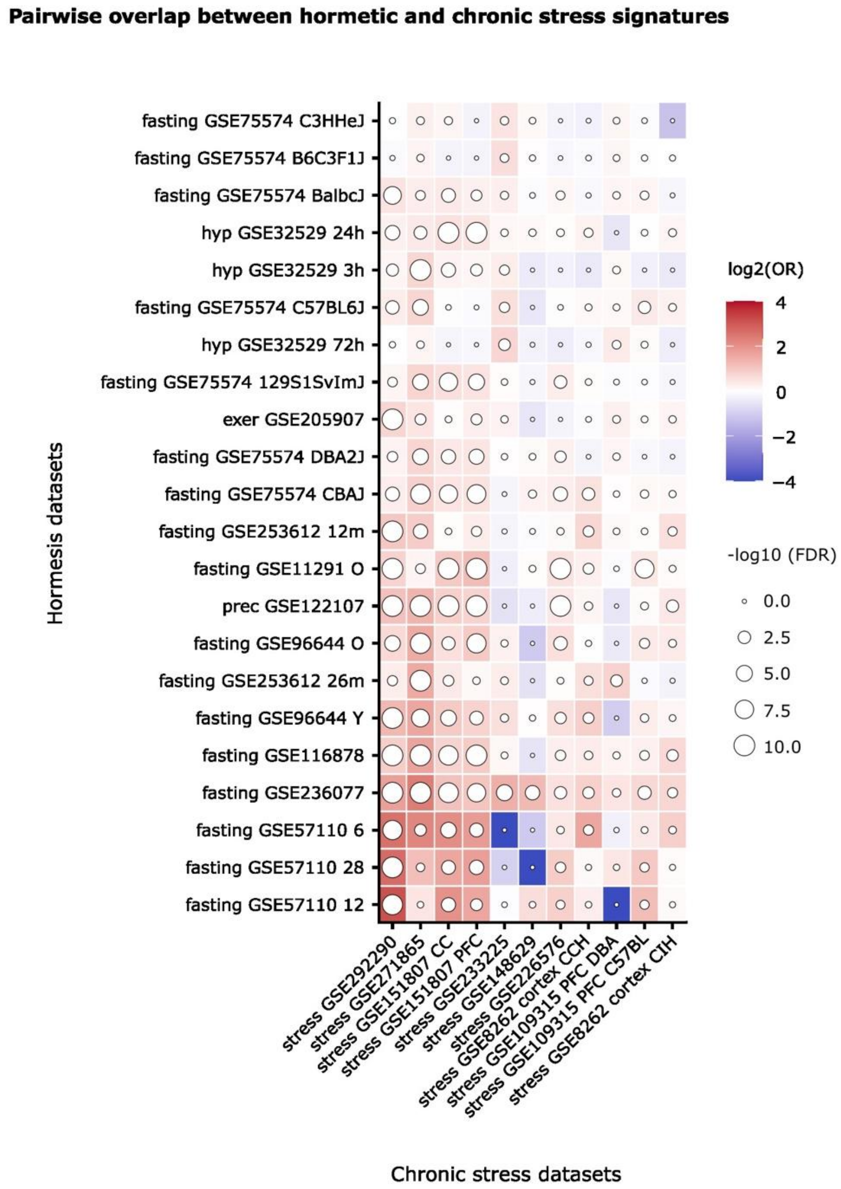


**Supplementary Figure 3. Substantial gene-level overlap between hormetic and chronic stress signatures.** Fisher's exact tests quantifying the overlap of differentially expressed genes between hormetic and chronic stress datasets. Circle size indicates false discovery rate (-$\log_{10}$(FDR)), while tile colour indicates the odd ratio (OR).

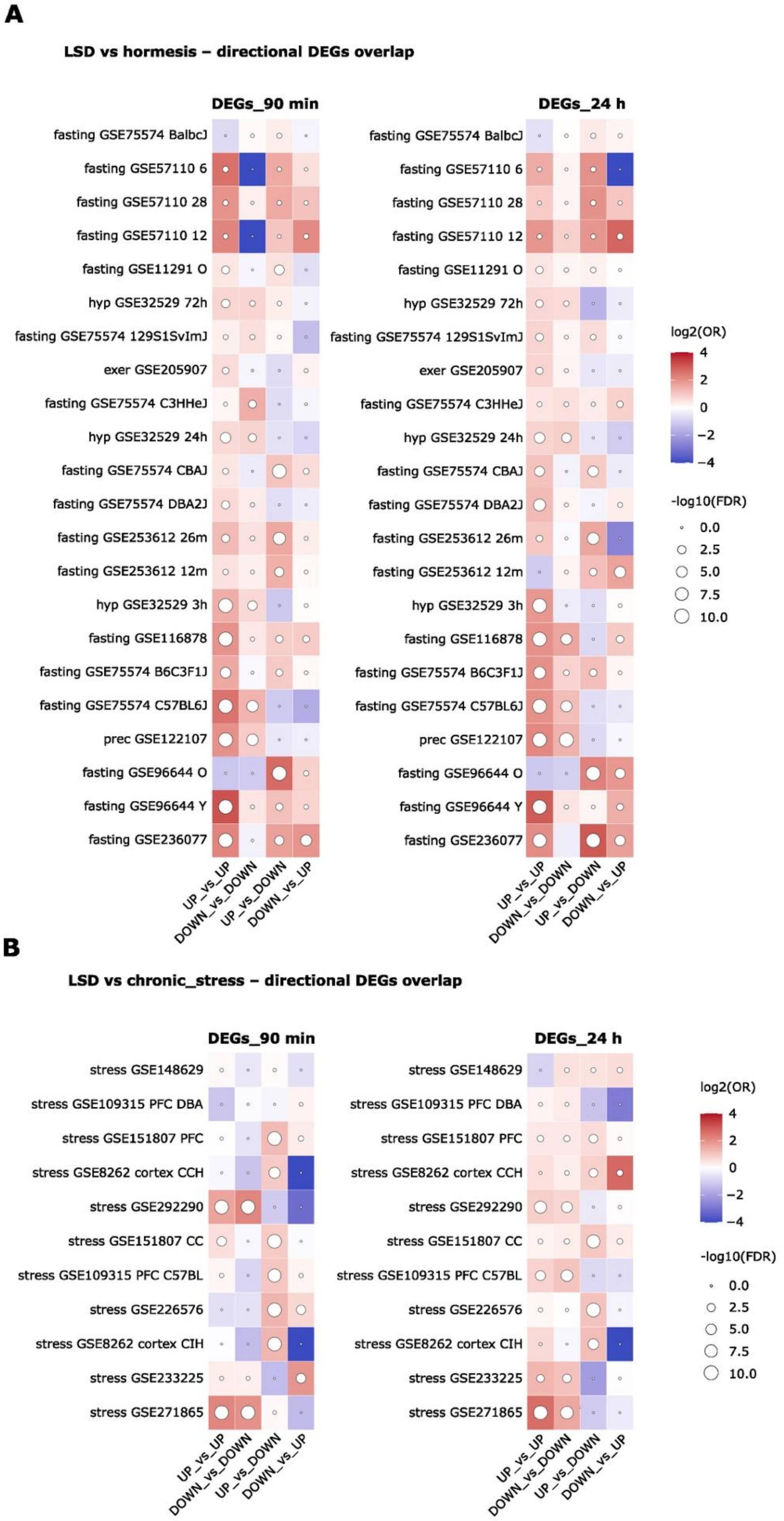


**Supplementary Figure 4. Direction-resolved overlap between LSD and stress-induced transcriptional programs. A)** Heatmaps showing direction-resolved overlap between differentially expressed genes (DEGs) induced by LSD vs hormetic stress paradigms, quantified using stratified Fisher's exact tests. Four comparisons were performed for each dataset: LSD_UP vs DATASET_UP, LSD_DOWN vs DATASET_DOWN, LSD_UP vs DATASET_DOWN, and LSD_DOWN vs

DATASET_UP. Results are shown separately for LSD signatures at 90 min (left) and 24 h (right). Circle size represents statistical significance (-$\log_{10}$ FDR), while tile colour encodes the odds ratio (OR). (**B**) Same analysis as in (**A**), applied to chronic stress datasets.

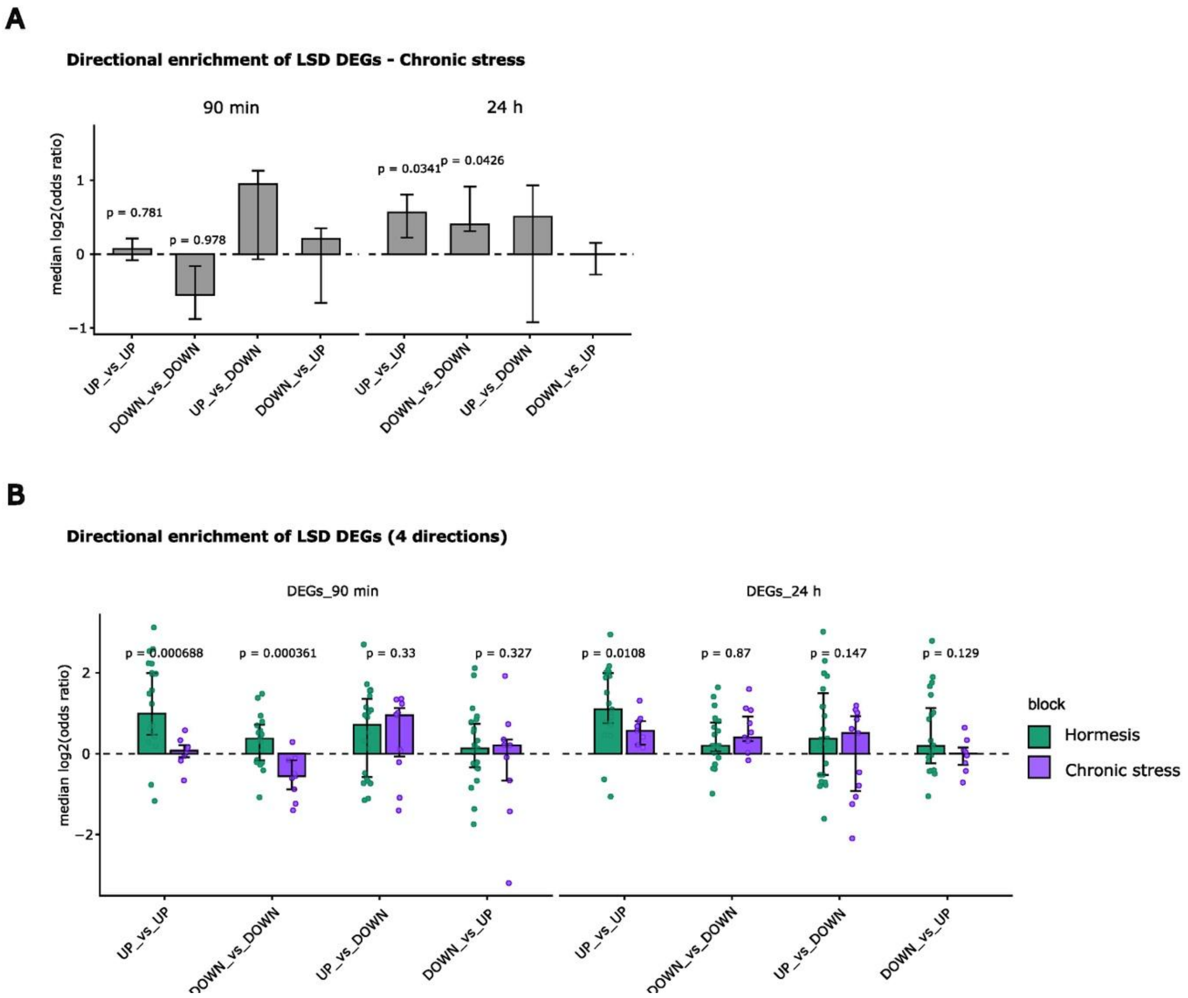


**Supplementary Figure 5. Direction-specific comparison of LSD overlap with hormetic and chronic stress paradigms. A)** Directional overlap between LSD and chronic stress transcriptional signatures. Odds ratios from direction-resolved Fisher's exact tests comparing LSD-induced DEGs with chronic stress signatures at 90 min and 24 h post-LSD (Fisher tests across LSD_UP vs DATASET_UP, LSD_DOWN vs DATASET_ DOWN, LSD_ UP vs DATASET_ DOWN, and LSD_ DOWN vs DATASET_UP). We tested whether ORs for UP_vs_UP or DOWN_vs_DOWN Fisher tests were higher than tests on discordant signs (UP_vs_DOWN and DOWN_vs_UP) via a one-sided Wilcoxon test. **B)** Direct comparison of odds ratios (ORs) obtained from stratified Fisher's exact tests for all four direction combinations between LSD-induced DEGs and external datasets. For each direction combination, ORs are shown separately for overlaps with hormetic (green columns) and chronic (purple columns) stress datasets at 90 min and 24 h post-LSD. Statistical significance was obtained via a one-sided Wilcoxon rank-sum test.

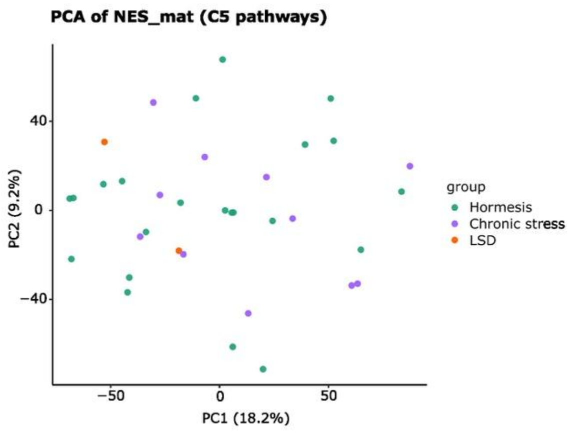


**Supplementary Figure 6. Pathway-level PCA enables cross-dataset integration of heterogeneous transcriptomic profiles.** PCA performed on pathway-level normalized enrichment score (NES) matrices across all datasets. Each point represents a dataset. The percentages indicate the variance explained by each component.

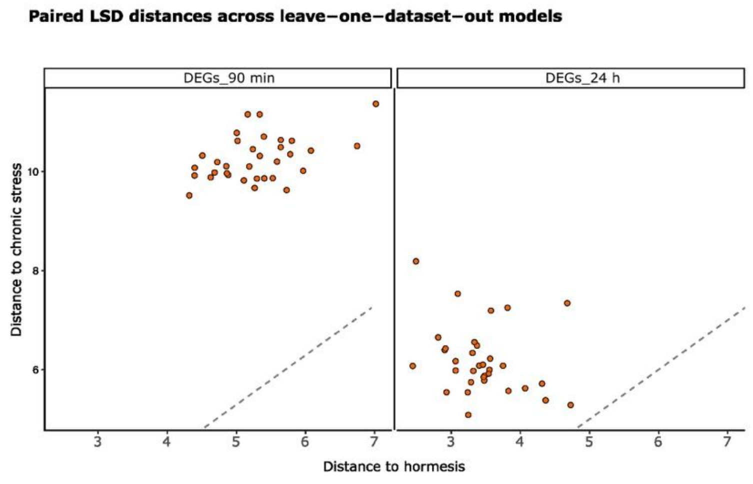


**Supplementary Figure 7. Leave-one-dataset-out stability of sPLS-DA-based positioning of LSD signatures.** Leave-one-dataset-out (LODO) analysis of the sPLS-DA model trained on hormetic and chronic stress datasets at the pathway level. Each point represents a single LODO iteration in which one training dataset was excluded, the model was refit, and LSD transcriptional signatures were projected into the resulting discriminant space. The figure shows the Euclidean distance between the LSD signatures and the hormesis centroid and chronic stress centroid for each refit model. Distances are reported separately for each LSD signature (90 min and 24 h). Lower values indicate greater proximity in the discriminant space.

**Supplementary Tables**

**Uploaded as additional files:**

**Supplementary Table 1** - List of stress datasets included in this study

**Supplementary Table 2** - Differentially Expressed Genes (DESeq2) at 90 min, 24 h and at both time points

**Supplementary Table 3** - Gene Ontology enrichment for up-regulated genes at 90 min, 24 h and at both time points

**Supplementary Table 4** - Top 20 categories with the highest loadings along sPLS-DA components
